# Modeling Continuous Admixture

**DOI:** 10.1101/033944

**Authors:** Ying Zhou, Hongxiang Qiu, Shuhua Xu

## Abstract

Human migration and human isolation serve as the driving forces of modern human civilization. Recent migrations of long isolated populations have resulted in genetically admixed populations. The history of population admixture is generally complex; however, understanding the admixture process is critical to both evolutionary and medical studies. Here, we utilized admixture induced linkage disequilibrium (LD) to infer occurrence of continuous admixture events, which is common for most existing admixed populations. Unlike previous studies, we expanded the typical continuous admixture model to a more general admixture scenario with isolation after a certain duration of continuous gene flow. Based on the extended models, we developed a method based on weighted LD to infer the admixture history considering continuous and complex demographic process of gene flow between populations. We evaluated the performance of the method by computer simulation and applied our method to real data analysis of a few well-known admixed populations.

## Introduction

Human migrations involve gene flow among previously isolated populations, resulting in the generations of admixed populations. In both evolutionary and medical studies of admixed populations, it is essential to understand admixture history and accurately estimate the time since population admixture because genetic architecture at both population and individual levels are determined by admixture history, especially the admixture time. However, the estimation of admixture time is largely dependent on the precision of the applied admixture models. Several methods have been developed to estimate admixture time based on the Hybrid Isolation (HI) model (Xu and Jin 2008; Price *et al*. 2009; Loh *et al*. 2013; Qin *et al*. 2015) or intermixture admixture model (IA) (Zhu *et al*. 2004), which assumes that the admixed population is formed by one wave of admixture at a certain time. However, the one-wave assumption often leads to under-estimation when the progress of the true admixture cannot be well modeled by the HI model. Jin et al. showed earlier that under the assumption of HI, the estimated time is half of the true time when the true model is a gradual admixture (GA) model (Jin *et al*. 2013).

Admixture models can be theoretically distinguished by comparing the length distribution of continuous ancestral tracts (CAT) (Gravel 2012; Jin *et al*. 2012; Ni *et al*. 2015), which refer to continuous haplotype tracts that were deviated from the same ancestral population. CAT inherently represents admixture history as it accumulates recombination events. Short CAT always indicates long admixture histories of the same admixture proportion, whereas long CAT may indicate a recent gene flow from the ancestral populations to which the CAT belongs. Based on the information it provides, CAT can be used to distinguish different admixture models and estimate corresponding admixture time. However, accurately estimating the length of CAT is often very difficult.

Weighted linkage disequilibrium (LD) is an alternative tool that can be used to infer admixture (Loh *et al*. 2013; Pickrell *et al*. 2014). Previous studies have indicated that this tool is more efficient than CAT because it requires neither ancestry information inference nor haplotype phasing, which often provides false recombination information, thus decreasing the power of estimation. Weighted LD has already been used in inferring multiple-wave admixtures (Pickrell *et al*. 2014; Zhou *et al*. 2015) However, these methods tend to summarize the admixture into different independent waves, even if the true admixture is continuous. In our previous work (Zhou *et al*. 2015), we mathematically described weighted LD under different continuous models, allowing us to determine admixture history using these models.

In the present study, we first developed a weighted LD-based method to infer admixture with HI, GA, and continuous gene flow (CGF) models (Pfaff *et al*. 2001), (Fig 1). Both GA and CGF models assume that gene flow is a continuous process. Next, we extended the GA and CGF models to the GA-I and CGF-I models, respectively (Fig 1), which model a scenario with a continuous gene flow duration followed by a period of isolation to present. We applied our method to a number of well-known admixed populations and provided information that would help better understanding the admixture history of these populations.

**Fig 1:**
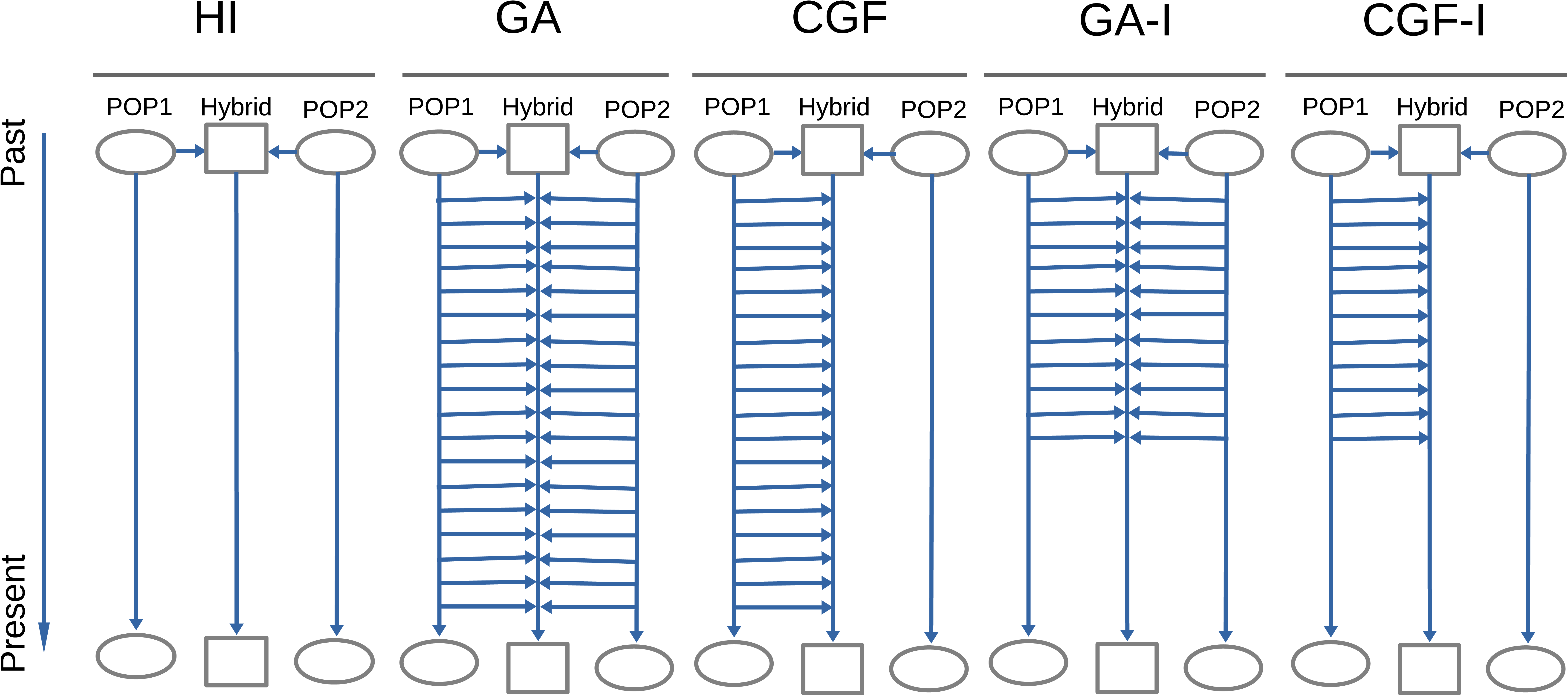
Classic admixture models (HI, GA and CGF) and the models we extended (GA-I and CGF-I). For each model, the simulated admixed population (Hybrid) is in the middle of two source populations (POP1 and POP2). Each horizontal arrow represents the direction of gene flow from the source populations to the admixed population. Once the genetic components flow into the admixed population, the admixed population randomly hybridizes with other existing components. The existence of horizontal arrows indicates gene flow from the corresponding source population.

## Material and Methods

### Datasets

Data for simulation and empirical analysis were obtained from three public resources: Human Genome Diversity Panel (HGDP) (Li *et al*. 2008), the International HapMap Project phase III (The International HapMap Consortium 2007) and the 1000 Genomes Project (1KG) (The 1000 Genomes Project Consortium 2012). Source populations for simulations are the haplotypes from 113 Utah residents with Northern and Western European ancestries from the CEPH collection (CEU) and the 113 Africans from Yoruba (YRI).

### Inferring Admixture Histories by using the HI, GA, and CGF Models

The expectation of weighted LD under a two-way admixture model has been described in detail in another work (Zhou *et al*. 2015). Following the previous notation, the expectation of weighted LD statistic between two sites separated by a distance *d* (in Morgan) is as follows:

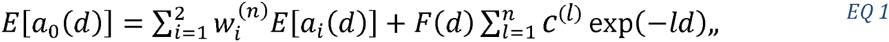
where 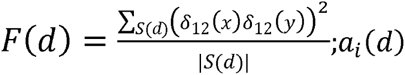, *i* = 0,1,2 are the weighted LD statistic of the admixed population (*i* = 0) and the source population *i*, (*i* = 1,2), respectively; *m_i_* is the admixture proportion from the source population *i*; and *δ*_12_(*x*) is the allele frequency difference between populations 1 and 2 at site *x;S*(*d*) is the set holding pairs of SNPs of distanced; *d;c*^(^*^l^*^)^ is admixture indicator for the admixture event of *l* generations ago, and *n* is supposed to be the number of generations ago when the source populations first met. To eliminate the confounding effect due to background LD from the source populations, we used the quantity, *z*(*d*), defined as follows, to represent the admixture induced LD (ALD) (Zhou *et al*. 2015).

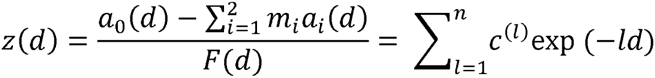

We presented it in a more compact form using the inner product of two vectors as follows:

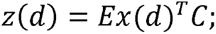
where

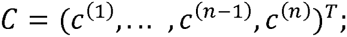
and

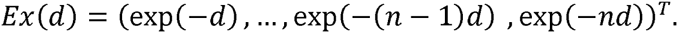

For different admixture models where admixture began *n* generations ago, z(d) varies in terms of the vector of coefficients of exponential functions (Zhou **et al**. 2015):

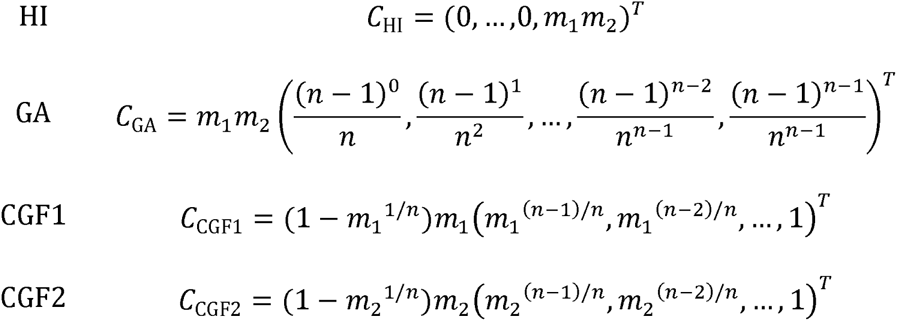
where the vector *C*_model_ has length *n* using the HI, GA, CGF1, or CGF2 model; and *n* represents when the admixture occurred (HI) or began (GA and CGF) in terms of generations. For different models, the coefficient vectors have different patterns (Fig 2), which can be used to infer the best-fit model for a certain admixed population.

**Fig 2:**
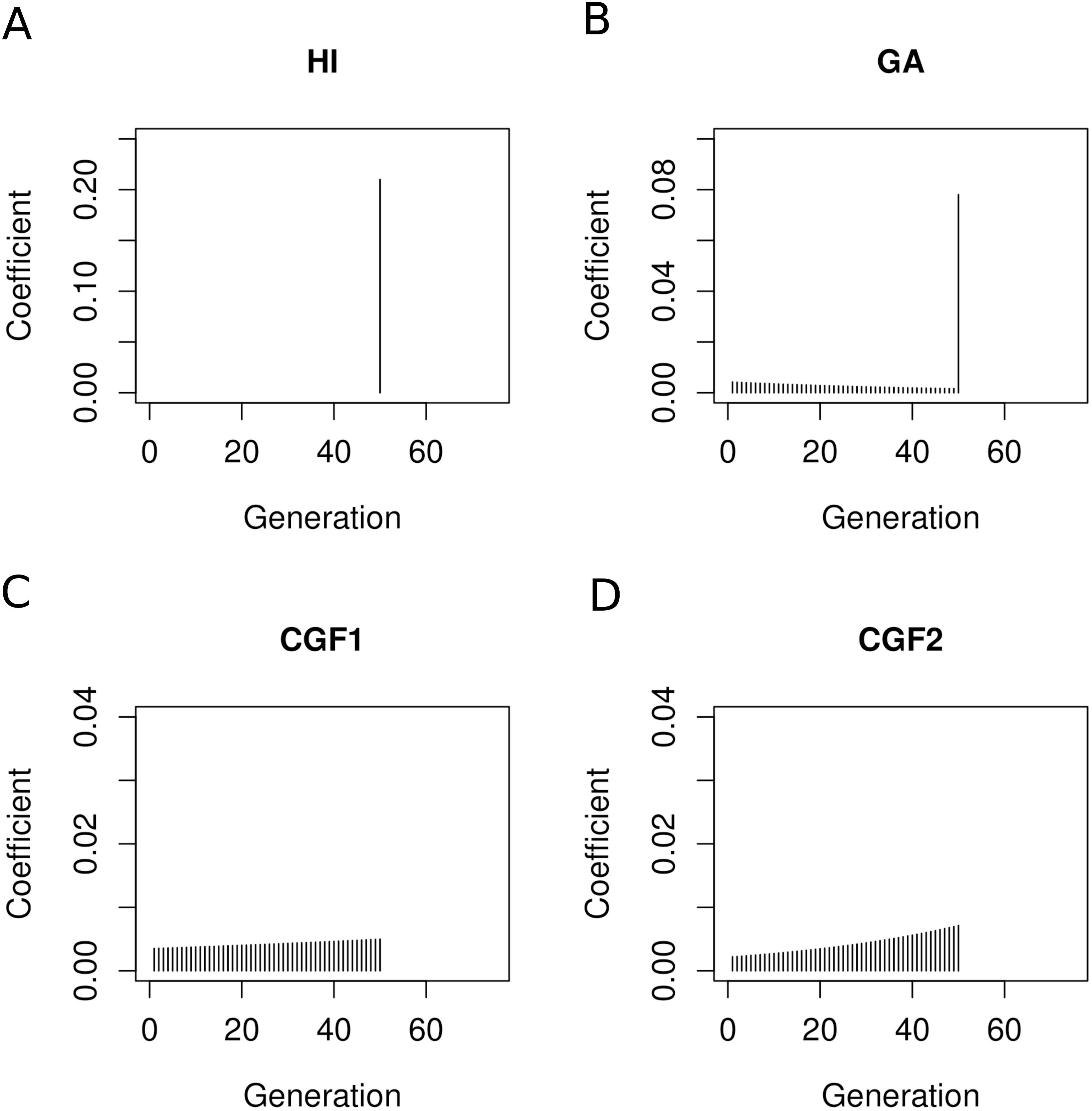
Coefficient vector of exponential functions for each model. For each admixture model, the starting time of the population admixture is 50 generations ago.

In the CGF model, CGF1 represents the admixture where source population 1 is the recipient of the gene flow from population 2, whereas CGF2 indicates source population 2 as gene flow recipient from population 1. Inference of the admixture time assuming the true admixture history is one of these different models that can be regarded as minimizing the objective function as follows:

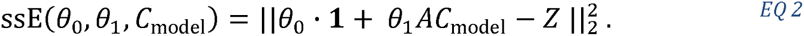

The optimization problem is therefore expressed as follows:

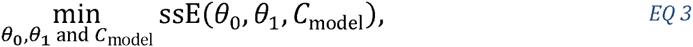
where *Z* = (*z*(*d*_1_), *z*(*d*_2_), …, *z*(*d_I_*))*^T^* is the observed ALD calculated from the single nucleotide polymorphism (SNP) data of both the parental populations and the admixed population; *θ*_0_ is a real number used to correct the population substructure; *θ*_1_ is a scalar that improves estimation robustness; **1** ∈ *R^I^* is a vector with each entry being 1; *A* is an *I* × *J* matrix with the *i*th row vector defined as *Ex*(*d_i_*)*^T^*, *i.e., A* = (*Ex*(*d*_1_), *Ex*(*d*_2_), …*, Ex*(*d_I_*))*^T^*, and *J* ≥ *n* is a pre-specified upper bound of *n*. Our definitions are consistent since we can let all entries be 0 after the *n*-th entry in *C*_model_.

Next, we tried to estimate the parameters *θ*_0_, *θ*_1_, and *C*_model_, where *C*_model_ has the information of the admixture model and the related admixture time *n* (in generations). In our analysis, the value of *n* is assumed to be a positive integer; therefore, our method is to go through all possible *n* values (with a reasonable upper limit *J*) to estimate *n* with the minimum value of the objective function. Given *n*, we used linear regression to estimate (*θ*_0_,*θ*_1_) such that the objective function was minimized. Using this approach, the value of *n* in relation to the minimal objective function value for each model was determined, which represents the time of admixture occurrence under each model.

### Admixture Inference under HI, GA-I, and CGF-I Models

GA and CGF models assume that the admixture is strictly continuous from the beginning of admixture to present. This assumption seems too strong to be valid in empirical studies. Here, we extended the GA model and CGF model to GA-I model and CGF-I model, respectively, by considering continuous admixture followed by isolation. In this case, the admixture event lasts from *G*_start_ generations ago to *G*_end_ generations ago. Similar to the previous case, the coefficients of exponential functions can be represented as the vector of length *G*_start_ for each model, whose first *G*_end_ − 1 entries are filled with zeros. Suppose the admixture lasted for *n* generations, then

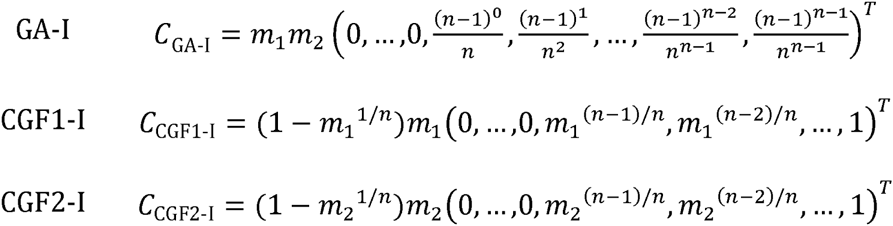

In this case, we can also try to find the parameters to minimize the objective function (EQ 2) under new models. By examining all possible pairs of (*G*_end_, *G*_start_), it is possible to determine the global minimum of the objective function, although this might not be computationally efficient Here, we used a faster algorithm (***Algorithm 1***) to determine the starting and ending time points of admixture.

Let *E* and *S* be the ending and starting time points (in generations, prior to the present) of the admixture, which we want to search for to minimize the objective function. The search starts from (*E*^0^, *S*^0^) = (1,*J*), where *J* is the upper bound for the beginning of the admixture event, which can be set to be a large integer to seek for a relatively ancient admixture event In our analysis of recent admixed populations, we set *J* = 500. For *k* = 1,2,…, (*E^k^,S^k^*) is updated from (*E^k^*^−1^,*S^k^*^−1^) by two alternative proposals. For convenience, we define

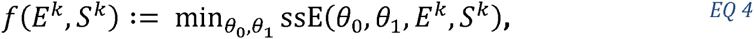
where *θ*_0_, *θ*_1_ can be determined by linear regression.

We choose the proposal that results in a smaller value for *f*. The search stops when the value of *f* with (*E^k^*^−1^,*S^k^*^−1^) is no larger than that of either proposal or *E^k^* = *S^k^*. In this way, we can readily estimate the time interval of the admixture event (*G*_end_, *G*_start_) quickly.

#### Algorithm 1

*for k in* 1,2,…

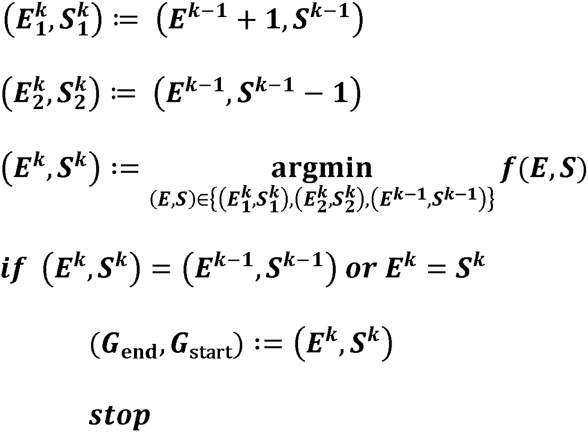

### Result evaluation

To check our assumption of the true history and evaluate the inference, an intuitive way is to compare empirical weighted LD with the fitted LD. Here, we use two quantities: msE and Quasi F, defined by the following:

1. Let *e* = *θ*_0_ · 1 + *θ*_1_*AC*_model_ − *Z*. We look at 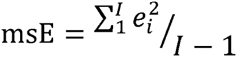 with *e_t_* being the *i*th entry of *e*. This reflects goodness of fit and strength of background noise. A smaller msE indicates less background noise and better fit.
2. Let 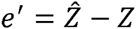, where 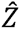 is the fitted weighted LD obtained from MALDmef, which theoretically can be regarded as the de-noised weighted LD. *e*′ is a vector of length *I*, with the *i*th entry denoted by 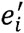. We look at the quasi-F statistic 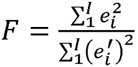. A small *F* indicates that the current fit does not significantly deviate from the previous fit.

A reliable result should have both small msE and small *F* values. Particularly, *F* is involved in model comparison: when *F* is too large, one would suspect that the true admixture history is far from any one of these models. Both *F* and msE are involved in revealing data quality. If *F* is small but msE is large, one would suspect that the quality of data is not good enough to draw convincing conclusions. Further explanation of these statistics is in Results and Discussion sessions.

### Identification of the best-fit model

For the convenience of illustration, we define the **core model** as the model used to infer admixture time. When inferring admixture of a target population, HI, GA, CGF1, CGF2, GA-I, CGF1-I and CGF2-I are used as the core models for conducting inference. Because GA-I, CGF1-I and CGF2-I describe more general admixture models than GA, CGF1, and CGF2, we classified model selection into two cases: one case is to identify the best-fit model(s) among the HI, GA, CGF1, and CGF2 models, whereas the more general case is to determine the best-fit model(s) among HI, GA-I, CGF1-I and CGF2-I models. In both cases, the same strategy is adopted, which depends on the pairwise paired difference of pseudo log(msE) values associated with each core model, which will be defined later. For an admixed population, there are *N* + 1 observed weighted LD curves obtained as follows: *N* (typically 22) autosomal chromosomes are considered in an individual genome, and one weighted LD curve is calculated from all these *n* chromosomes while the other *N* weighted LD curves are obtained by jackknife resampling, leaving out one chromosome for each LD curve (Loh *et al*. 2013; Pickrell *et al*. 2014; Zhou *et al*. 2015). Next, we fit each observed weighted LD curve for each core model by estimating *θ*_0_,*θ*_1_ and the time interval, which in turn allowed us to obtain the msE value associated with the optimal parameters for each weighted LD curve. Taken together, a total of *N* + 1 msE values associated with *N* + 1 LD curves were evaluated in each core model. For model *M*, the log(msE) obtained from all *N* chromosomes is denoted by 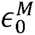 and that from the LD curve with the *q*-th chromosome left out by 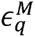. Following Tukey (Tukey 1958), we defined the *q*-th pseudo log(msE) for model *M* to be 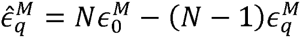 and treated these pseudo values approximately as independent Next, we defined the best-fit core model(s) to be the model(s) with significantly small 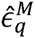. A pairwise Wilcoxon signed-rank test was conducted for the pseudo log(msE) of the four models. More precisely, Wilcoxon signed-rank test is applied to all pairs of models with the 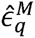 being paired by index *q*, and then the p-values are adjusted to control familywise error rate (Table 1). We used the Holm-Bonfferroni method to adjust p-values (Holm 1979). When 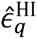 were not significantly larger than those of the best model, i.e., the model associated with the smallest sample median of pseudo log(msE) values, HI was selected. Otherwise the models whose 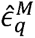 were not significantly larger than those of the best model were selected (the best model was selected as well). The significance level was set to be 0.05. Here, we paired the pseudo values according to index *q* and used Wilcoxon signed-rank test on the paired differences because according to our experience, 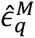 are strongly correlated with *q* and hence *q* is a major covariate that must be controlled in the test to gain higher power. This is also why even though theoretically there are examples where the best model according to our definition can be significantly worse than another model in our process, we believe such extreme cases are unlikely in practice and still use this method. In addition, log(msE) rather than msE were used because after logarithm transformation small values of msE can also have large effect to the comparison. That is, we could better detect the difference between small msE, thus gaining greater power in the test. This claim is also justified by our experience.

**Table 1:** Adjusted p-values of pairwise Wilcoxon signed-rank test among core models: HI, GA-I, CGF1-I, CGF2-I.

| True Model | Best Model(s) | Adjusted p-Values of Pairwise Wilcoxon Signed Rank Test |  |  |  |  |  |
| --- | --- | --- | --- | --- | --- | --- | --- |
|  |  | HI: GA-I | HI: CGF1-I | HI: CGF2-I | GA-I: CGF1-I | GA-I: CGF2-I | CGF2-I: CGF1-I |
| **HI (100)** | HI | 0.97 | 0.14 | 0.97 | 0.012 | 0.15 | 0.97 |
| **HI (50)** | HI | 0.98 | 0.98 | 0.52 | 0.70 | 0.16 | 0.16 |
| **CGF1 (1-100)** | CGF1-I | 0.012 | 0.012 | 0.012 | 0.012 | 0.012 | 0.012 |
| **CGF1 (1-50)** | CGF1-I, CGF2-I | 0.012 | 0.012 | 0.012 | 0.041 | 0.064 | 0.055 |
| **GA (1-100)** | GA-I | 0.012 | 0.012 | 0.012 | 0.012 | 0.012 | 0.012 |
| **GA (1-50)** | GA-I | 0.012 | 0.012 | 0.012 | 0.012 | 0.020 | 0.012 |
| **CGF1-I (30-100)** | HI | 0.19 | 0.55 | 0.15 | 0.012 | 0.55 | 0.012 |
| **CGF1-I (70-100)** | HI | 0.97 | 0.97 | 0.97 | 0.020 | 0.012 | 0.20 |
| **GA-I (30-100)** | CGF1-I, GA-I | 0.012 | 0.012 | 0.012 | 0.30 | 0.012 | 0.020 |
| **GA-I (70-100)** | HI | 0.52 | 0.52 | 0.52 | 0.029 | 0.012 | 1 |

In each column, the adjusted p-values of the Wilcoxon signed-rank test comparing the two models are presented for all simulation cases. Simulated true model is followed by the parenthesis of time interval for the corresponding gene flow, where the first term in the parenthesis is the ending time of the admixture and the second term is the beginning time of the admixture. They are in the measurements of generation before present. For HI model, only one time point is included in the parenthesis.

### Software

Our algorithm has been implemented in an R package (R Core Team 2014), named CAMer (Continuous Admixture Modeler). The package is available on the website of population genetic group: http://www.picb.ac.cn/PGG/resource.php or on Github: https://www.github.com/david940408/CAMer.

## Results

### Simulation studies

Admixed populations were simulated in a forward-time way under different admixture models with the software **AdmixSim** (Yang 2015). Simulation was initiated with the haplotypes from source populations (YRI and CEU) and haplotypes for the admixed population were generated by resampling haplotypes with recombination from source populations and the admixed population of last generation. During the simulation, population size was kept as 5000 and migration rates was controlled by the admixture model with the final admixture proportion in the admixed population to be 0.3. We employed a mono recombination map in our simulation, which means recombination rate between two markers is positively proportional to their physical distance. For each model, simulation was performed using 10 replicates; each replicate contained 10 chromosomes with a total length of 3 Morgans. To evaluate the performance of our algorithm, we simulated admixed populations under the following conditions:

1. HI of 50 and 100 generations, designated as HI (50) and HI (100),
2. GA of 50 and 100 generations, designated as GA (1-50) and GA (1-100), respectively,
3. CGF of 50 and 100 generation, population 1 as the recipient, designated as CGF1 (1-50) and CGF1 (1-100) respectively,
4. CGF-I of a 70-generation admixture followed by 30-generation isolation, and a 30-generation admixture followed by a 70-generation isolation, with population 1 as the recipient, designated as CGF1-I (30-100) and CGF1-I (70-100) respectively, and,
5. GA-I of a 70-generation admixture followed by a 30-generation isolation and a 30-generation admixture followed by a 70-generation isolation, designated as GA-I (30-100) and GA-I (70-100), respectively.

With simulated admixed populations, we first used the HI, GA and CGF models as core models to conduct inference (Fig S1). When the simulated model was a HI, GA, or CGF model, our method was able to accurately estimate the simulated admixture time, as well as to determine the correct model, with an accuracy of 73.33%. When the simulated model was a CGF-I or GA-I model, the estimated time based on the core model HI was within the time interval of the admixture, whereas all best-fit models were HI (Table 2). This result has indicated the limitation of using the GA and CGF models in inferring admixture history.

**Table 2:**
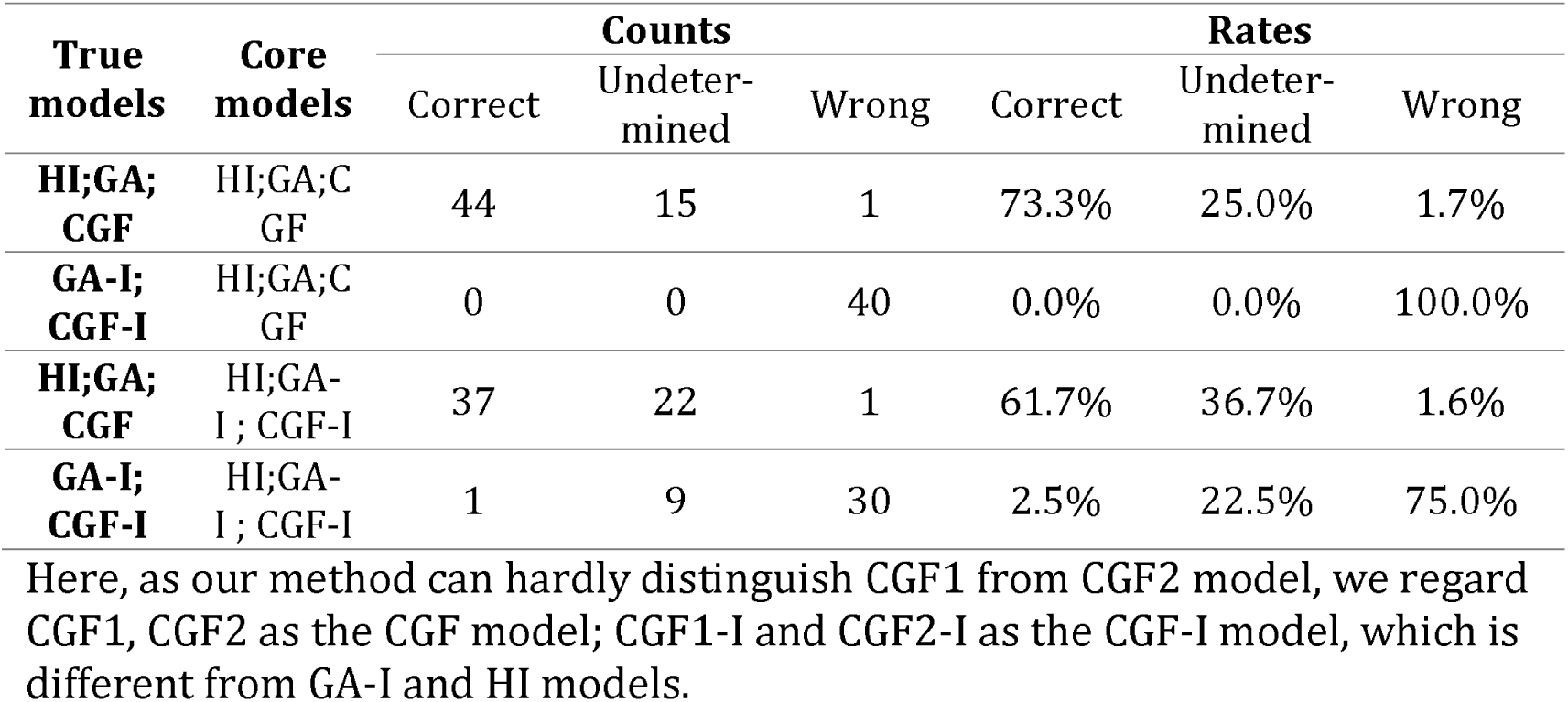
Accuracy of model detection

| True models | Core models | Counts |  |  | Rates |  |  |
| --- | --- | --- | --- | --- | --- | --- | --- |
|  |  | Correct | Undetermined | Wrong | Correct | Undetermined | Wrong |
| **HI;GA;CGF** | HI;GA;CGF | 44 | 15 | 1 | 73.3% | 25.0% | 1.7% |
| **GA-I;CGF-I** | HI;GA;CGF | 0 | 0 | 40 | 0.0% | 0.0% | 100.0% |
| **HI;GA;CGF** | HI;GA-I;CGF-I | 37 | 22 | 1 | 61.7% | 36.7% | 1.6% |
| **GA-I;CGF-I** | HI;GA-I;CGF-I | 1 | 9 | 30 | 2.5% | 22.5% | 75.0% |

2 Here, as our method can hardly distinguish CGF1 from CGF2 model, we regard CGF1, CGF2 as the CGF model; CGF1-I and CGF2-I as the CGF-I model, which is different from GA-I and HI models.

Using the same simulated admixed populations, we then employed GA-I, CGF-I and HI as core models for performing inference (Figs 3 and S2-S11). With HI, GA, or CGF considered as the true model, our estimation of the optimal model remained highly accurate. On the other hand, when the true model was GA-I or CGF-I, the failure rate decreased by 25%, compared to the estimation in the previous setting. Furthermore, the estimated time intervals were wider than those of the true ones, although the findings were still more accurate than those using GA and CGF as core models (Table 2).

**Fig 3:**
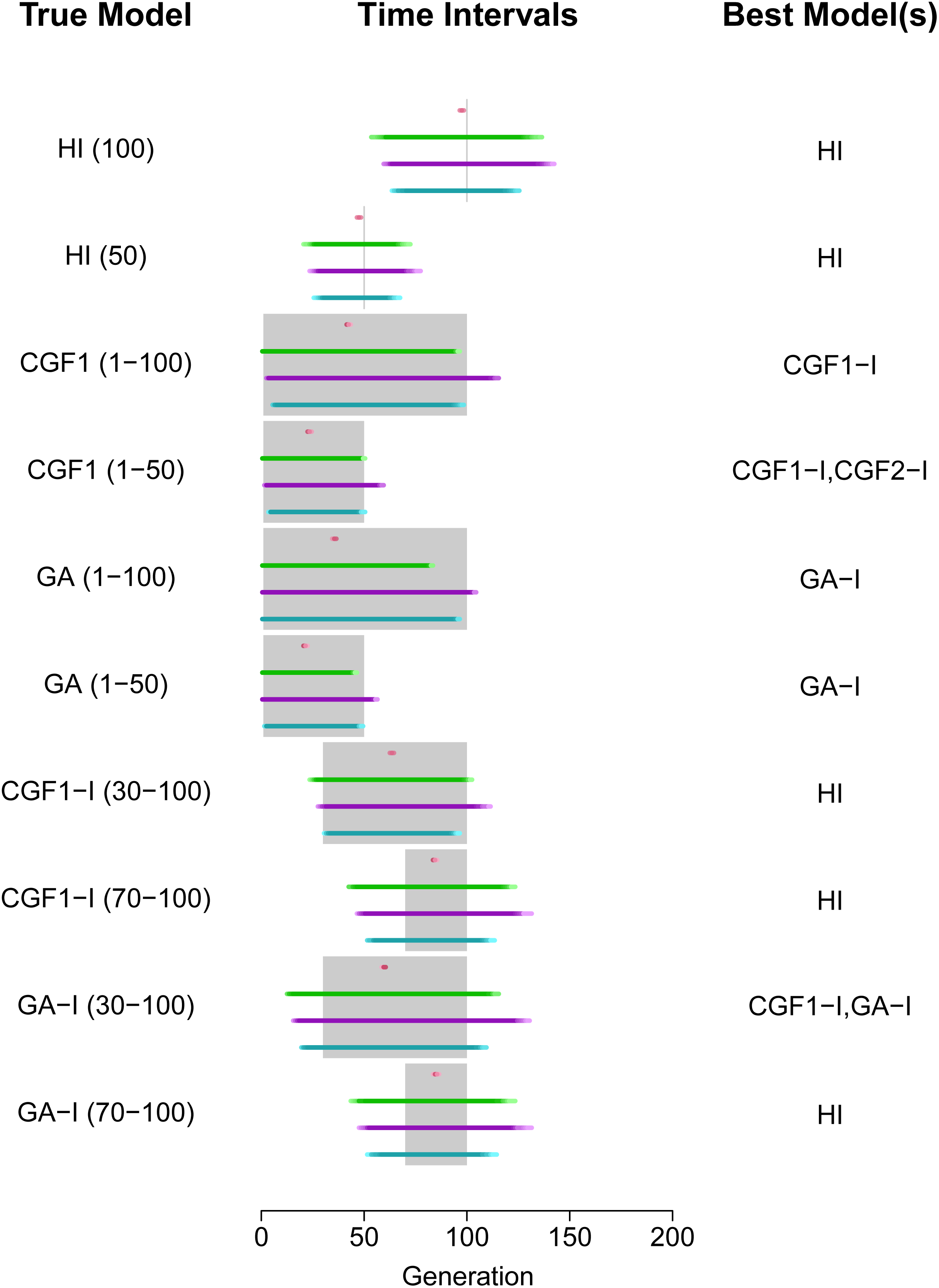
Evaluation of CAMer under various simulated admixture models. Here, the core models are HI, GA-I, CGF1-I, and CGF2-I. The simulated models (True Model) are listed on the left, with the admixture time interval depicted in the parentheses. The gray area on the middle vertical panel is the simulated time interval, whereas colored lines indicate the estimated time intervals under different core models. HI: pink; CGF1-I: green; CGF2-I: purple; GA-I: blue. The intensity of lines means the number each point is covered by the time intervals estimated from all jackknives. Lighter colors represent fewer covers while darker colors mean more.

### Empirical analysis

We applied CAMer to the selected admixed populations from HapMap, HGDP, and 1KG. For each target population, we first used MALDmef to calculate the weighted LD and fit the weighted LD with hundreds of exponential functions (Zhou *et al*. 2015). Next, with the weighted LD of target populations, we determined the admixture model and estimated admixture time with CAMer. Quasi F and msE are designed for evaluating the inference with CAMer. The value of msE usually indicates data quality: small msE may indicate a high signal-to-noise ratio (SNR) and vice versa. The quasi F value measures the goodness of fit of the model we employed to fit the admixture event. A small *F* value indicates that the model we used was of satisfactory performance in modeling an admixture event. In our analysis, we used 10^−5^ as the threshold for msE and 1.5 as the threshold for *F*. Therefore, when the msE value ≤ 10^−5^ and the *F* value ≤ 1.5, we could not “reject the null hypothesis” that the related model was the true model, i.e., the model well fit the admixture event. On the other hand, an msE value ≥ 10^−5^ indicates low-quality data that is incapable of identifying the best-fit model, whereas an *F* value ≥ 1.5 prompts us to “reject the null hypothesis” and concludes that the model did not well fit the admixture. In the case of the same population from different databases, the data with smaller msE values were given more credits. For example, we obtained samples of ASW from the HapMap and the 1KG. With the ASW data (CEU and YRI as source populations) from HapMap, the best-fit model was HI of 6 generations, and both msE and F values indicated that the inference was acceptable (Fig S12). Similarly, using the ASW data (CEU and YRI as source populations) from 1KG, the best-fit model was HI of 6 generations (Fig S13). However, a quasi F value of 2.54 indicated that HI model did not satisfactorily fit the admixture event. Because the msE value of the data set from 1KG was smaller, the conclusion using ASW was as follows: based on the best data we had, the time intervals estimated under the HI, GA-I, CGF1-I, and CGF2-I model were 6 generations, 1–9 generations, 1–13 generations, and 1–9 generations, respectively. Furthermore, none of these models satisfactorily modeled the admixture, whereas the HI model showed better performance. We also applied CAMer to other admixed populations (Table 3, Figs S14–17). MEX (source poulations: CEU (64 individuals) and American Indian (7 Colombians, 14 Karitiana, 21 Maya, 14 Pimas and 8 Suruis)) was satisfactorily modeled by the CGF1 model or GA-I model, with the estimated admixture time interval being 1–17 or 2–16 generations, respectively. We also analyzed Eurasian populations, which showed that the Uygurs (source populations: Han (n = 34) and French (n = 28)) most likely fit a continuous model, with a gene flow lasting for more than 60 generations to the present or near present. We cannot determine which model fits best. However, the values of msE were all larger than 10^−5^, indicating that the results were not so reliable. The Hazara population (source populations: Han (n = 34) and French (n = 28)) experienced a GA-I-like admixture event that lasted for about 58 generations, which started 63 generations ago and ended approximately 5 generations ago.

**Table 3:**
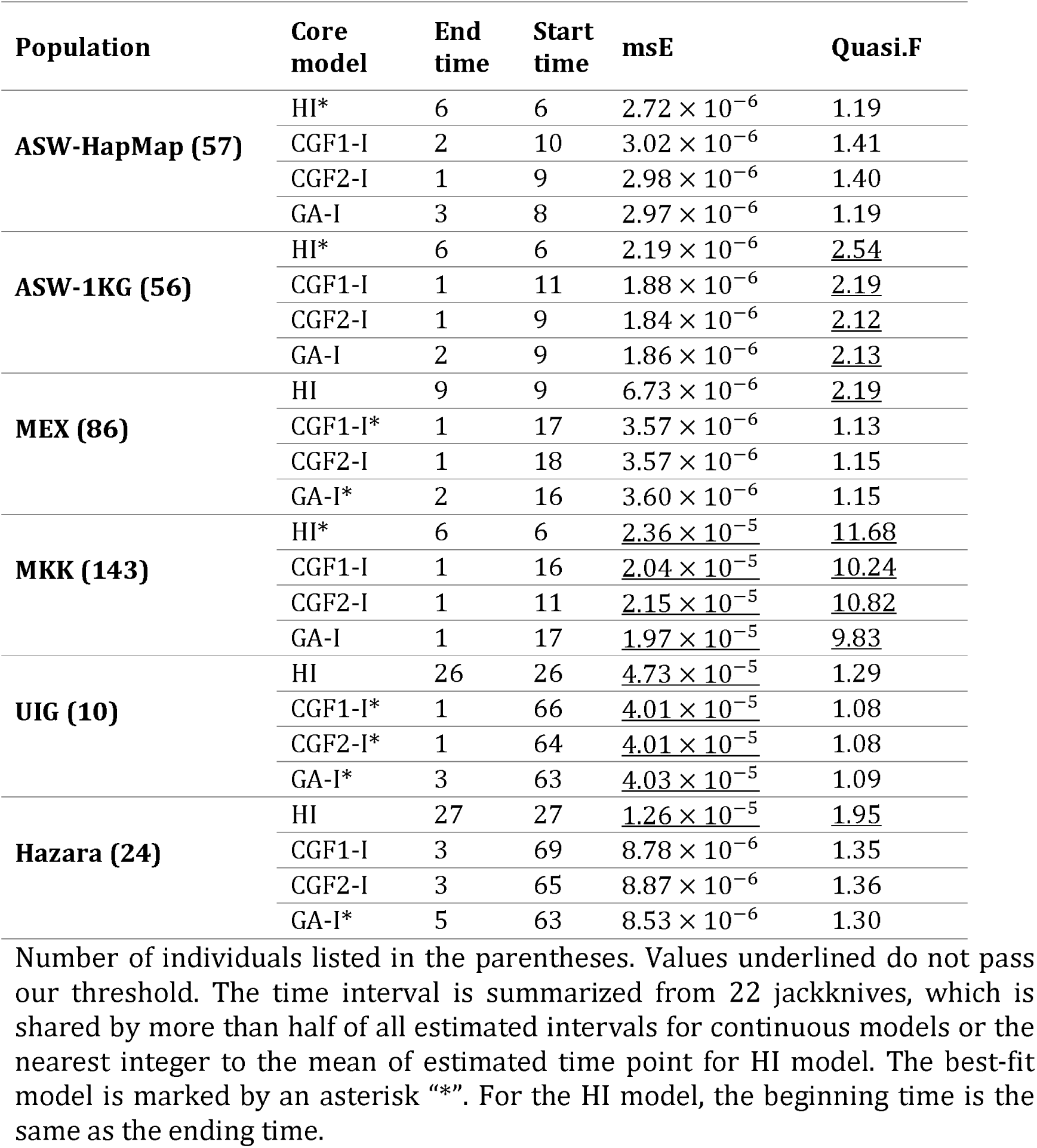
Results of CAMer on empirical populations

| Population | Core model | End time | Start time | msE | Quasi.F |
| --- | --- | --- | --- | --- | --- |
| **ASW-HapMap (57)** | HI* | 6 | 6 | $2.72 \times 10^{-6}$ | 1.19 |
| | CGF1-I | 2 | 10 | $3.02 \times 10^{-6}$ | 1.41 |
| | CGF2-I | 1 | 9 | $2.98 \times 10^{-6}$ | 1.40 |
| | GA-I | 3 | 8 | $2.97 \times 10^{-6}$ | 1.19 |
| **ASW-1KG (56)** | HI* | 6 | 6 | $2.19 \times 10^{-6}$ | <u>2.54</u> |
| | CGF1-I | 1 | 11 | $1.88 \times 10^{-6}$ | <u>2.19</u> |
| | CGF2-I | 1 | 9 | $1.84 \times 10^{-6}$ | <u>2.12</u> |
| | GA-I | 2 | 9 | $1.86 \times 10^{-6}$ | <u>2.13</u> |
| **MEX (86)** | HI | 9 | 9 | $6.73 \times 10^{-6}$ | <u>2.19</u> |
| | CGF1-I* | 1 | 17 | $3.57 \times 10^{-6}$ | 1.13 |
| | CGF2-I | 1 | 18 | $3.57 \times 10^{-6}$ | 1.15 |
| | GA-I* | 2 | 16 | $3.60 \times 10^{-6}$ | 1.15 |
| **MKK (143)** | HI* | 6 | 6 | $2.36 \times 10^{-5}$ | <u>11.68</u> |
| | CGF1-I | 1 | 16 | $2.04 \times 10^{-5}$ | <u>10.24</u> |
| | CGF2-I | 1 | 11 | $2.15 \times 10^{-5}$ | <u>10.82</u> |
| | GA-I | 1 | 17 | $1.97 \times 10^{-5}$ | <u>9.83</u> |
| **UIG (10)** | HI | 26 | 26 | $4.73 \times 10^{-5}$ | 1.29 |
| | CGF1-I* | 1 | 66 | $4.01 \times 10^{-5}$ | 1.08 |
| | CGF2-I* | 1 | 64 | $4.01 \times 10^{-5}$ | 1.08 |
| | GA-I* | 3 | 63 | $4.03 \times 10^{-5}$ | 1.09 |
| **Hazara (24)** | HI | 27 | 27 | $1.26 \times 10^{-5}$ | <u>1.95</u> |
| | CGF1-I | 3 | 69 | $8.78 \times 10^{-6}$ | 1.35 |
| | CGF2-I | 3 | 65 | $8.87 \times 10^{-6}$ | 1.36 |
| | GA-I* | 5 | 63 | $8.53 \times 10^{-6}$ | 1.30 |

3 Number of individuals listed in the parentheses. Values underlined do not pass 4 our threshold. The time interval is summarized from 22 jackknives, which is 5 shared by more than half of all estimated intervals for continuous models or the 6 nearest integer to the mean of estimated time point for HI model. The best-fit 7 model is marked by an asterisk “\*”. For the HI model, the beginning time is the 8 same as the ending time.

## Discussion

Modeling the demographic history of an admixed population and estimating time points of this particular event are essential components of evolutionary and medical research studies (Zhu *et al*. 2004; Zhu and Cooper 2007; Gravel 2012; Jin *et al*. 2012, 2013; Ni *et al*. 2015; Zhou *et al*. 2015). Previous methods have employed the length distribution of ancestral tracts (Gravel 2012; Jin *et al*. 2012, 2013), which highly depends on the result of local ancestral inference and haplotype phasing. Another limitation of earlier methods is that only HI, GA, and CGF models were utilized to fit the admixture as well as in identifying the best-fit model. In the present study, our simulations showed that when the true model was not HI, GA, or CGF, the generated inferences were relatively difficult to interpret.

Our method, CAMer, can be utilized in inferring admixture histories by using weighted LD, which can be calculated using genotype data with MALDmef (Zhou *et al*. 2015). Furthermore, we extended the GA and CGF models to the GA-I and CGF-I models in order to infer the time interval for a period of continuous admixture events followed by isolation. Although HI model is a degenerate case for both GA-I and CGF-I models, where the admixture window becomes 1 generation, we kept it in our method because it is the most popular model employed in previous admixture studies. Considering the difficulty in the fitting problem with exponential functions, it is in our expectation that CAMer was not consistently very accurate in determining the admixture model based on the weighted LD decay. However, its natural advantage of independence of both haplotype phasing and local ancestry inference makes it privilege to other CAT based method. And our simulations indicated that its time interval estimations were reliable when its assumption that the true admixture history could be well approximated by one of the core models is valid.

Two quantities, namely msE and quasi F, were used to check the assumption of our method stated above and evaluate the credibility of the models’ inference. These two quantities should both be taken into consideration to identify whether the models well describe the admixture history. Both the data quality and the goodness of fitting of models can affect the value of msE, although the *F* value mainly measures the goodness of modeling. Informally, for the convenience of interpretation, msE is considered to reveal the data quality and *F* value is considered to check model assumption on admixture history. In our analysis, we suggested thresholds for msE and *F* to determine whether the null hypothesis should be rejected or not, which may be too strict in empirical analysis. Actually, msE and *F* values together measure whether the observed weighted LD can be well fit by the best-fit model(s). For example, the fitting process showed poor performance in the MKK population, which was accompanied by exaggerated msE and *F* values, showing significant inconsistencies between the observed and fitted weight LD curves, which indicates that the true admixture history cannot be well explained by any of the core models (Fig S17). Therefore, in empirical analysis, one can informally think that the msE value reflects the quality of the data, whereas *F* value describes the performance of the model, although both of them measure the goodness of fitting.

In our previous study (Zhou *et al*. 2015), we fit the weighted LD with hundreds of exponential functions. However, this approach did not fully reveal the occurrence of continuous admixture. To address this issue, the present study developed CAMer to model admixture as a continuous process. CAMer also employed extensions of the classic continuous models, GA-I and CGF-I, which may bring the bias to have a wider admixture window when the real admixture exists in a short time. But it is still proved to be able to give more credible estimations in modeling population admixture.

Taken together, CAMer is a powerful method to model a continuous population admixture, which in turn would help us elucidate the complex demographic history of population admixture.

## Author contributions

Conceived and designed the study: **SX**. Developed methods and computer tools: **YZ HQ**. Analyzed the data: **YZ** and **HQ**. Interpreted the data and wrote the paper: **SX YZ HQ**.

## Acknowledgements

None.

